# Pyruvate from bone marrow mesenchymal stem cells supports myeloma redox homeostasis and anabolism

**DOI:** 10.1101/2024.08.08.607157

**Authors:** Elías Vera-Sigüenza, Cristina Escribano-Gonzalez, Irene Serrano-Gonzalo, Kattri-Liis Eskla, Charlotte Speakman, Alejandro Huerta-Uribe, Lisa Vettore, Himani Rana, Adam Boufersaoui, Hans Vellama, Ramin Nashebi, Ielyaas Cloete, Jennie Roberts, Supratik Basu, Mark Drayson, Christopher Bunce, Guy Pratt, Fabian Spill, Oliver D.K. Maddocks, Daniel A. Tennant

## Abstract

Multiple myeloma is an incurable cancer of plasma cells that depends on the bone marrow for its survival. Despite its prevalence, the molecular mechanisms underlying this malignancy remain poorly understood. In this study, we aim to bridge this knowledge gap by elucidating the metabolic interplay between myeloma cells and bone marrow mesenchymal stem cells (BMMSCs). BMMSCs are crucial in supporting myeloma cell metabolism, contributing to their proliferation, survival, and resistance to chemotherapy. Through a combination of mathematical modelling and experimental co-cultures, we demonstrate that pyruvate – the end product of glycolysis – plays a key role in myeloma cell metabolism. Our findings reveal that myeloma cells predominantly rely on the uptake of pyruvate produced by neighbouring BMM-SCs via the plasma membrane proton-linked monocarboxylate transporters MCT-1 and MCT-2 encoded by the Slc16a1 and a2 genes, respectively. Furthermore, we show that pharmacological inhibition of the MCT-1/2, with AZD3965, triggers a cascade of compensatory metabolic responses, disrupting redox balance and significantly reducing the proliferation capacity of co-cultured myeloma cells.

## 1. Introduction

Multiple myeloma (MM) is a haematological cancer characterised by antibody-producing malignant plasma cells in the bone marrow (BM). Unlike other blood malignancies, MM typically has a low circulating tumour burden, as malignant plasma cells rarely circulate en masse, unlike in plasma cell leukaemia [1, 2, 3, 4, 5].

The myeloma tumour microenvironment (TME) plays a crucial role in supporting the malignancy’s proliferation, potentially contributing to the disease’s poor prognosis [4, 5, 6]. Factors within the bone marrow environment, including metabolic interactions, cytokines, and direct cell-to-cell contact, have been described as essential for this support [7, 8, 9, 10, 11, 12, 13]. Similar interactions have been observed in other cancers, such as colon cancer and pancreatic ductal adenocarcinoma, where malignant cells rely on stromal cells to enhance the tumour microenvironment through redox-sensitive metabolic networks. Examples of this include previously described stromal-cancer cell subnetworks that exchange pyruvate for lactate or pyruvate for alanine for survival [11, 14, 15, 16]. In multiple myeloma, the major cell type thought to support the malignant plasma cell through these multiple pathways is the bone marrow mesenchymal stem cell (BMMSC) [8, 9, 10, 11, 17, 18].

It is becoming increasingly clear that monocultured cells in vitro are insufficient to allow the modelling of the metabolic network found within the TME [3, 19, 20, 21]. Recent advancements by us and others using in vitro co-cultures, primary cells, and mathematical modelling of the myeloma TME have shed new light on the malignant bone marrow [9, 19, 20, 22, 23, 24]. Employing a bespoke constraint-based model (CBM) of the myeloma-bone marrow mesenchymal stem cell dyad, we computationally and experimentally demonstrated the formation of a metabolic consortium within the multiple myeloma TME [24]. We hypothesised that this consortium leads to profound metabolic reprogramming in both co-cultured myeloma cells and BMM-SCs [24, 25].

In this study, we further integrate computational modelling with experimental work to demonstrate that metabolic shuttling from bone marrow mesenchymal stem cells (BMMSCs) to malignant plasma cells vitally maintains the redox balance in myeloma and enhances their proliferation [6, 24, 26, 27]. Using both immortalised and primary autologous myeloma and bone marrow mesenchymal stem cells, in co-culture, we establish that a major trafficked metabolite is pyruvate, which contrasts with previously hypothesised movement of lactate [9, 10, 22]. Consequently, we show that BMMSC-derived pyruvate serves as a crucial alternative carbon source for maintaining the redox potential of myeloma cells. Finally, we explore the mechanisms and wider metabolic consequences of this dependency. We carry out pharmacological inhibition of pyruvate import by targeting monocarboxylate transporter 1 & 2 (MCT-1/MCT-2). We demonstrate that reducing exogenous pyruvate import to myeloma cells leads to reduced proliferation and a maladaptive metabolic network. The results presented here pinpoint pyruvate and the monocarboxylate transporters 1 & 2 (MCT-1/MCT-2) as promising metabolic targets for novel therapeutic interventions in the fight against this disease [28, 29, 30, 31, 32].

## 2. Materials and Methods

### Cell cultures

HS-5 and JJN-3 cell lines, purchased from ATCC, were maintained in RPMI-1640 (Sigma-Aldrich, R8758) with 10% foetal bovine serum (FBS - Sigma-Aldrich, F7524) at 37*^◦^*C in a humidified incubator with 5% CO_2_ [33, 34]. Cells were tested for mycoplasma contamination every 6 months by PCR. Passaging of adherent stromal cells, or HS-5 cell line, was performed using 0.05% Trypsin (Gibco, 12-605-036). Treatment with AZD3965 (Stratech Scientific, S7339-SEL) was at 10 *µ*M where indicated, with dimethyl sulfoxide (DMSO) as vehicle control [28]. For co-culture experiments, HS-5 cells were seeded at 12 *×* 10^4^ cells per well of a 6-well plate. Next day, JJN-3 were seeded on top of a trans-well (Appleton Woods, CC401) at 8 *×* 10^4^ cells per well.

### Primary patient samples

Patients diagnosed with multiple myeloma (MM) and monoclonal gammopathy of undetermined significance (MGUS) were recruited from a specialist clinic at Haematology, University Hospitals Birmingham NHS Foundation Trust, Queen Elizabeth (QE) Hospital Birmingham, UK. The study, reference 14-178, received approval from the local ethics committees at South Birmingham, Birmingham East, North, and Solihull, and informed written consent was obtained in accordance with the Declaration of Helsinki in all cases.

For patient samples with multiple myeloma and monoclonal gammopathy of undetermined significance, bone marrow was diluted in Red Blood Cell Lysis Solution (Miltenyi Biotec, 130-094-183) according to the manufacturer protocol. Cell pellet was then resuspended in RPMI-1640, cells counted, spun down for washing and resuspended in RPMI-1640 supplemented with 1% ITS+ (Corning, 11593560), 1% Penicillin/Streptomycin (Merck, P0781) and 1% Amphotericin B (Merck, A2942). BMMSCs were defined by adherence to culture flasks and myeloma cells remained in suspension or were lightly adherent to the BMMSCs. Media was replaced every 5 to 7 days in order to maintain co-cultures. To facilitate stable isotope tracing experiments, antibiotics were removed a week before. Patient number and characteristics are shown in Appendix A.

When required, myeloma patient cells were obtained using StraightFrom Whole Blood and Bone Marrow CD138 MicroBeads as per manufacturer’s instructions (130-105-961, Miltenyi Biotec) for positive selection from PBMCs from blood fraction separation with a density gradient (Histopaque 1119 and 1077, Sigma-Aldrich).

### Quantitative Real-Time PCR

RNA from cell lines was extracted using the RNeasy Mini-Kit in accordance with the manufacturer’s protocol (Qiagen, UK). RNA from patient samples was extracted using the Total RNA Purification Micro Kit in accordance with the manufacturer’s protocol (Norgen Biotek, Canada). 1 *µ*g RNA was transcribed using the Reverse Transcription System kit (Promega, UK). Quantitative PCR (qPCR) was performed on QuantStudio5 Real-Time PCR System using the TaqMan gene expression master mix (Applied Biosystems, UK) [35]. The following probes were used: Slc16a1 (MCT-1, Hs01560299_m1_), Slc16a7 (MCT-2, Hs00940851_m1_) and Slc16a3 (MCT-4, Hs00358829_m1_); Thermo Fisher Scientific, UK. Gene expression levels were normalised to ACTB (Actin, HS01060665_g1_). Relative expression levels were calculated using the 2-ΔΔCT method and compared to JJN-3 - as outlined by Schmittgen and Livak [36].

### Glucose measurements

Media was collected from each well, cells were spun down and supernatant was taken for glucose analysis. Levels were measured using a Contour XT glucometer (Bayer).

### Western blotting

Cells were washed once (PBS, pH 7.4) before scraping into Laemmli buffer (Sigma, 2301-1VL). Immunoblotting was then performed as previously described by Westbrook et al. [37]. After blocking, nitrocellulose membranes were incubated with MCT-1 (1:1000, Santa Cruz, sc-365501 for mouse samples; 1:1000, ProteinTech, 20139-1-AP for human samples), MCT-2 (1:1000, LabNed, LN0315312), MCT-4 (1:1000, Santa Cruz, sc-376140) and/or actin (1:4000 dilution, Sigma, A4700). After washing anti-rabbit-HRP (Cell Signalling, 7074S) or anti-mouse-HRP (Cell Signalling, 7076S) secondary antibodies were incubated with the membrane before binding was visualised using EZ-ECL (Biological Industries, 20-500-120) and imaging on a ChemiDoc Imaging System (BioRad) [38].

### Metabolic tracing

For tracing experiments, HS-5 and JJN-3 cell lines were plated to be 70% confluent after 48 hours. Media was then changed to flux media (modified RPMI-1640 without glucose, glutamine, L-isoleucine, L-leucine, L-valine, and phenol red, Cell Culture Technologies), with 11 mM glucose and 2 mM glutamine supplemented either in unlabelled form or as ^1^3C_6_-glucose and ^1^3C_5_-glutamine (CK Isotopes) where indicated with RPMI SILAC (Gibco, A2494201) or human-like plasma medium (HPLM - Gibco), as indicated. After 48 hours, 100 *µ*L of media was removed for extraction and analysis. JJN-3 cells were pelleted by centrifugation as above. The remaining media was aspirated and the HS-5 and cells or empty wells were washed twice with ice cold saline, after which 500 *µ*L MeOH was added. HS-5 cells were scraped and transferred to a cold Eppendorf. 500 *µ*L D_6_-glutaric acid in ice-cold water (1 *µ*g/mL) was then added (CDN isotopes, D-5227) and 500 *µ*L of chloroform (pre-chilled to −20*^◦^*C). After shaking on ice for 15 minutes and centrifugation, the polar phase was transferred to another tube for derivatisation, which was dried with vacuum centrifugation at 45*^◦^*C.

### Sample Extraction and NEM-Derivatisation for GSH Detection

HS-5 cells were seeded the day before to allow them to attach and reach approximately 70% confluency by the time of extraction. The next morning, the medium was replaced with flux medium, and JJN-3 cells were added on top of a transwell, supplemented with 2.5 mM GSH where indicated, for 6 hours of incubation.

After 6 hours, JJN-3 suspension cells were pelleted at 4*^◦^*C. The medium was removed, and the cell pellets were washed with a cold NEM-saline mixture (0.9% saline and 30% 20 mM NEM solution in high purity water). After removal of the NEM-saline solution, the pellets were lysed with 100 *µ*L of cold extraction solution (20% 20 mM NEM, 40% acetonitrile, 40% methanol, and 2 *µ*g/mL D_6_-glutaric acid).

For HS-5 adherent cells, the medium was removed and kept on ice. The cells were then washed with NEM-saline mixture, lysed with 100 *µ*L of extraction solution, and scraped off the plate on ice.

For medium extraction, samples were spun down to pellet cell debris. 20 *µ*L of the supernatant were taken, and 80 µL of medium extraction solution (6 mM NEM, 40% acetonitrile, 40% methanol, 2 *µ*g/mL D_6_-glutaric acid) were added. Finally, all samples were briefly vortexed, then spun down at high speed at 4*^◦^*C for 5 minutes. Supernatants were aspirated and transferred to glass vials ready for injection.

### Stable Isotope Tracing by LC-QTOF

2-10 *µ*l of sample were injected into an Agilent 1290 Infinity II Liquid Chromatography system containing an Atlantis Premier BEH Z-HILIC 1.7*µ*m 2.1*×*150mm Column (Waters) equipped with a Vanguard precolumn.

Solvent A was 20 mM Ammonium Bicarbonate (LiChropur, Sigma) with 0.1% v/v Ammonium Hydroxide (Alfa Aesor) in 90% Water (Honeywell)/10% Acetonitrile (BioSolv) containing 5 *µ*M Infinitylabs Deactivator Solution (Agilent). Solvent B was 90% Acetonitrile / 10% Water containing 5 *µ*M Infinitylabs Deactivator Solution. A non-linear binary gradient was used as follows: 0 min 85% B, 2 min 85%, 18 min 60% B, 22 min 30% B, 22.1 min 10% B, 25 min 10% B, 25.1 min 85% B, 30 min 85% B.

The column compartment was maintained at 30*^◦^* C throughout. Blanks were run every 6-9 samples to monitor for metabolite carry over. An inline Agilent 6546 Q-TOF was used for metabolite measurements operating in high resolution mode in negative electrospray ionisation. Mass calibration was performed immediately prior to each sample batch. The drying gas was 8l/min at 225*^◦^* degrees.

The nebuliser was at 30psi with a sheath gas temperature of 300 degrees at 12l/min. The VCap was 2000V with 500V nozzle voltage, fragmentor 125V, Skimmer 60V, Octapole RF 750V. MS1 scans were obtained at 1Hz between 50-1050 m/z. Continuously infused reference masses of 112.985587 and 1033.988109 provided real time mass correction throughout all analytical runs.

Raw data (D) was imported into Agilent MassHunter Profinder followed by batch isotopologue extraction against an inhouse Personal Compound Database and Library (PCDL) containing metabolite IDs with confirmed retention times. Glutathione was detected as it’s N-Ethylmaleimide derivative (GSH-NEM). To account for the incomplete mass resolution of dual elemental tracer (15N1,13C2) enriched GSH-NEM, mass tolerances on isotopologue extraction were relaxed to 30ppm to give a single M+3 peak. All peak integrations were manually reviewed and compared to unlabelled controls. Peak areas were subsequently normalised to an internal standard (2H6-glutaric acid) and cell count using inhouse scripts in Excel (Microsoft).

### LC-MS metabolomics

Sample analysis was conducted using an LC-MS platform comprising an Accela 600 LC system and an Exactive mass spectrometer (Thermo Scientific). Metabolite separation was achieved using a Sequant ZIC-pHILIC column (4.6 mm *×* 150 mm, 3.5 mm) (Merck), 20 mM ammonium carbonate in water (as mobile phase A) and 100% acetonitrile (as mobile phase B). Gradient elution started at 20% A, followed by a linear increase to 80% A at minute 30, 92% A for 5 min, and 20% A for 10 min, at flow rate of 300 *µ*L/min. Column temperature was set to 28*^◦^*C. The Exactive mass spectrometer was operated in full scan mode over a mass range of 70–1200 m/z at a resolution of 50,000 with polarity switching. Thermo raw files were converted into mzML files using the MSConvert tool (ProteoWizard) and imported to MZMine 2.3 for peak picking and sample alignment. An in-house made database was used for metabolite identification. Finally, peak areas were used for comparative quantification.

### Derivatisation and Gas Chromatography – Mass Spectrometry

Dried down extracts were derivatised using a two-step protocol. Samples were first treated with 2% methoxamine in pyridine (40 *µ*L, or 20 *µ*L if primary samples, 1h at 60*^◦^*C), followed by addition of N-(tert-butyldimethylsilyl)-N-methyl-trifluoroacetamide, with 1% tert-butyldimethylchlorosilan (60 *µ*L, or 30 *µ*L if primary samples, 1h at 60 *^◦^*C). Samples were transferred to glass vials for GC-MS analysis using an Agilent 8890 GC and 5977B MSD system. Sample injection, GC-MS conditions and metabolite identification was performed as previously described by Westbrook et al. [37]. GC-MS data were analysed using Agilent Mass Hunter software for real time analysis of data quality before conversion to .CDF format and analysis with in-house MATLAB scripts.

### Normalisation and Quantification

Cells were counted using counting chambers (Fast Read 102, Kova International). GCMS data were analysed using Agilent Mass Hunter software for real time analysis of data quality before conversion to .CDF format and analysis with in-house MATLAB scripts. Graphs and statistical analysis were performed using GraphPad Prism 9 and MATLAB.

### Mathematical Model and **^13^**C-Metabolic Flux Analysis

We developed and validated two genome-scale cell-specific *in-silico* constraint based models (CBMs) founded on the Recon3D framework [39, 24]. The construction, validation processes, and source code are extensively described in Vera-Siguenza et al. [24]. These models were intricately linked to replicate the co-culture conditions of our experiments, utilising the COBRA (Constraint-Based Reconstruction and Analysis) Toolbox along with a suite of algorithmic approaches. Source code for all algorithms and models which we employ here and in Vera-Siguenza et al. [24] can be freely accessed and obtained from our GitHub repository under an MIT open-source license [24, 39, 40].

To partially quantify and parameterise the metabolic fluxes of our constraint-based models, we employed ^13^C metabolic flux analysis (^13^C-MFA). Briefly, this method deduces intracellular flux patterns from mass isotopomer distributions measured via mass spectrometry. The process and established protocols were followed as described in Vera-Siguenza et al. [24], Antoniewicz [41], Young [42], and Rahim et al. [43]. We conducted this analysis using the ‘Isotopomer Network Compartmental Analysis’ (INCA) MATLAB routine, suitable for both steady-state and isotopically non-stationary metabolic flux analysis [42, 43].

We recalibrated our models with the data presented in the results section of this study - using the methodologies of Vera-Siguenza et al. [24], Antoniewicz [41], Young [42], and Rahim et al. [43]. Furthermore, we verified isotopic steady states within our co-culture system and confirmed that isotopic enrichment changes remained within established measurement boundaries. Our simulations incorporated data on ^13^C_6_-glucose and ^13^C_5_-glutamine. Estimations of glucose and glutamine uptake fluxes were approximately 0.45 *µ*mol/10^6^ cells/hr and 0.63 *µ*mol/10^6^ cells/hr, respectively, consistent with Antoniewicz [41] guidelines. We ensured the reliability of our model by conducting over 100 simulation restarts from diverse initial conditions. Flux results were analysed using chi-square (*χ*^2^) to assess goodness-of-fit, with 95% confidence intervals calculated for each flux estimate, as detailed in Vera-Siguenza et al. [24].

## 3. Results

### Pyruvate utilisation underpins enhanced metabolic efficiency in myeloma

Cell-cell interactions within the TME may be a critical factor in the development and survival of cancer [44, 45, 46]. Recent studies across various tumour types indicate that metabolites are trafficked between cancer cells and stromal cells to support the malignancy’s metabolic needs [13, 47, 48, 49]. To determine if comparable dynamics occur in multiple myeloma, we investigated glucose uptake as an indicator of carbon metabolism in both mono-culture and co-culture of the HS-5 and JJN-3 cell lines, representing bone marrow mesenchymal stem- and myeloma cells, respectively [12, 17]. In co-culture, our experiments showed that both HS-5 and JJN-3 cells consume less glucose per cell than when cultured individually (Supplementary Fig. 1), suggesting a more bioenergentically efficient glucose utilisation, and perhaps altered substrate preference. This observation aligns with the predictions of our CBM (Fig. 1a - see Materials and Methods section) and underscores the metabolic efficiency facilitated by the resulting metabolic network spanning both cell types [24]. This has been demonstrated in other malignancies like pancreatic and colon cancers, as well as myeloma [10, 11, 13, 17, 24, 49].

**Figure 1:**
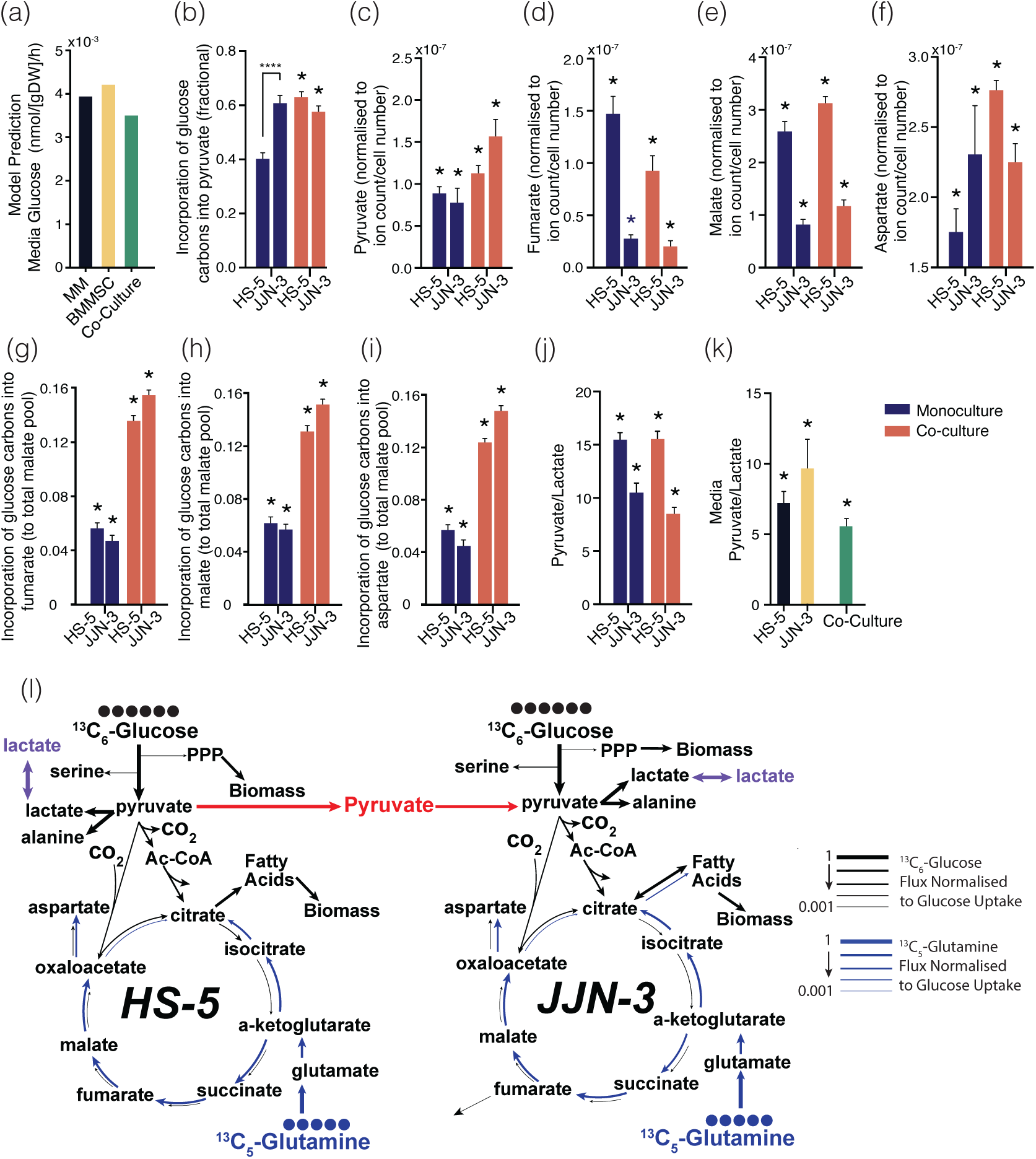
Metabolic adaptations in monoculture versus co-culture of HS-5 and JJN-3 cells. Data are displayed as ave. ± S.D. *, p*<*0.05, two-tailed student t-test. All experiments were repeated 3 times. **(a).** Metabolic utilisation in co-culture as predicted by constraint-based modelling (CBM), with both cell types consuming less glucose per cell compared to monoculture. For a comparison with experimental data see Vera-Siguenza et al. [24]. **(b).** Fractional enrichment from glucose to pyruvate in JJN-3 cells under both conditions, contrasting with increased pyruvate production in HS-5 cells when co-cultured. (***p*<*0.001, ****p*<*0.0001 from ANOVA and Dunn’s multiple comparison) **(c).** Variations in pyruvate levels in HS-5 cells in co-culture, while JJN-3 cells maintain consistent levels. **(d-f).** Consistent m+2 labelling of TCA cycle intermediates malate and fumarate in both cell types, with a marked increase in aspartate labelling in co-cultured JJN-3 cells. **(g).** Normalised incorporation of glucose-derived carbons into the fumarate pool for both cell types, indicating similar uptake in each condition. **(h).** Glucose carbons incorporated into malate, with comparable levels in monoculture and co-culture. **(i).** Increased incorporation of glucose-derived carbons into the aspartate pool in JJN-3 cells during co-culture. **(j).** Intracellular pyruvate/lactate ratio of both cell types. **(k).** Pyruvate/lactate ratio in the media is significantly reduced in co-culture compared to monoculture, suggesting altered metabolic interactions between HS-5 and JJN-3 cells when cultured together. **(l).**Metabolic flux map obtained from ^13^C-MFA simulations of HS-5 and JJN-3 cells in co-culture. Arrow widths are proportional to flux magnitudes. For numerical values, refer to the Supplementary Material.

We therefore decided to investigate this using ^13^C_6_-glucose both in monoculture and co-culture conditions. We found that JJN-3s demonstrated no significant change in the fractional enrichment of pyruvate from glucose. In contrast, HS-5s exhibited a marked increase in this process when co-cultured with JJN-3s, when compared with monoculture (Fig. 1b). HS-5s had notable pyruvate level changes when co-cultured with JJN-3s cells, whereas JJN-3s maintained similar pyruvate levels in both conditions (Fig. 1c). Unexpectedly, the conversion of pyruvate to lactate in HS-5s showed no significant differences between monoculture and co-culture (Supplementary Fig. 2).

We also noted consistent labelling of malate and fumarate in both HS-5 and JJN-3 cells. Both of which are important compounds in the tricarboxylic acid (TCA) cycle. A notable distinction was seen in co-cultured myeloma (JJN-3) cells, which exhibited a significant increase in aspartate labelling, a pattern not mirrored by HS-5 cells (Figs.1d - 1f). This observation is particularly relevant given aspartate’s critical role as a limiting factor in cell proliferation [50, 51, 52, 53, 54]. Overall, our findings indicate that the incorporation of glucose-derived label into malate, fumarate, and aspartate (m+2 isotopologues) was markedly higher in co-culture compared to monoculture. The result suggests either enhanced pyruvate contribution to TCA cycle metabolites in both cell types when in co-culture or may reflect metabolite exchange between cell types (Figs.1g - 1h and Supplementary Fig. 3).

It has previously been suggested that in other cell co-culture systems, metabolite movement between stromal and cancer cells may facilitate the maintenance of a redox state in the cancer cells that could support a less constrained metabolic network. We therefore examined the lactate to pyruvate ratio in both cell types and the medium in both monoculture and co-culture conditions – often used as a surrogate for the cytosolic NAD^+^/NADH ratio. Interestingly, the intracellular ratio was observed to remain similar in both cell types between monoculture and co-culture, albeit lower in the HS-5 cell lines (Fig. 1j), suggesting that the two cell types maintain different cytosolic redox states regardless of culture conditions. However, when we examined the extracellular media from both cell types in monoculture, we observed that the media from JJN-3 cells was markedly different from their intracellular ratio, whereas the media from HS-5 cells was comparable. This suggests that the flux of pyruvate and lactate across the plasma membrane in JJN-3 cells may not be equimolar (Fig. 1k). The ratio of these metabolites in co-culture was further lowered compared to in monoculture, and remained similar to that of the HS-5 cell. These results suggested that the two cell types demonstrate differential flux of pyruvate and lactate across their plasma membranes, and that in co-culture, the exchange of these metabolites may be used by the cells to maintain favourable redox homeostasis.

To investigate the changes in metabolism in co-culture in more detail, we conducted parallel ^13^C_6_-glucose and ^13^C_5_-glutamine metabolic flux analyses (data stored in Supplementary Material - GitHub). Through these simulations, we observed a predicted increase in the directional flux of pyruvate from HS-5 to JJN-3 cells, supporting the change in extracellular lactate/pyruvate ratio observed experimentally (Fig. 1l - red). In the same simulation, a portion of these carbons entered the myeloma mitochondria to undergo further oxidation via the TCA cycle. Here, we observed 40% formed citrate and around 18% continued to form *α*-ketoglutarate. Our modelling suggests a predicted citrate transfer from the mitochondria to the cytoplasm in myeloma cells (Fig. 1l - blue). Given the nitrogen requirement of the pyruvate-alanine transamination predicted, it was not surprising that our model confirmed previous studies suggesting that glutamine is a crucial nutrient in myeloma, with its carbons contributing to the citrate pool (Fig. 1l) [24, 55, 52, 56, 57, 58].

The model also predicted that while pyruvate flux may occur between cell types, this was in the absence of overall flux of lactate. This is a departure from its previously assumed role in myeloma redox balance (Fig. 1l - purple) but in agreement with results previously suggested by our CBM [9, 10, 22, 24].

### MCT-1 transporter activity supports the metabolic relationship between BMMSCs and myeloma cells

Pyruvate and lactate are transported between cell types through the action of monocarboxylate transporters, which have varying affinities for these two substrates [59]. To examine which of these transporters were expressed by HS5 and JJN3 cells, we assessed the expression of monocarboxylate transporters previously reported to have the highest affinity for pyruvate and/or lactate: 1, 2 and 4 (encoded by Slc16a1, Slc16a7 and Slc16a3, respectively). While MCT-1 and 2 were expressed in both cell lines (HS-5s and JJN-3s), as well as a wider panel of myeloma cell lines (human U266 and mouse 5TGM1 cells), MCT-4 was not detected in the JJN-3 cell line (Fig. 2a).

**Figure 2:**
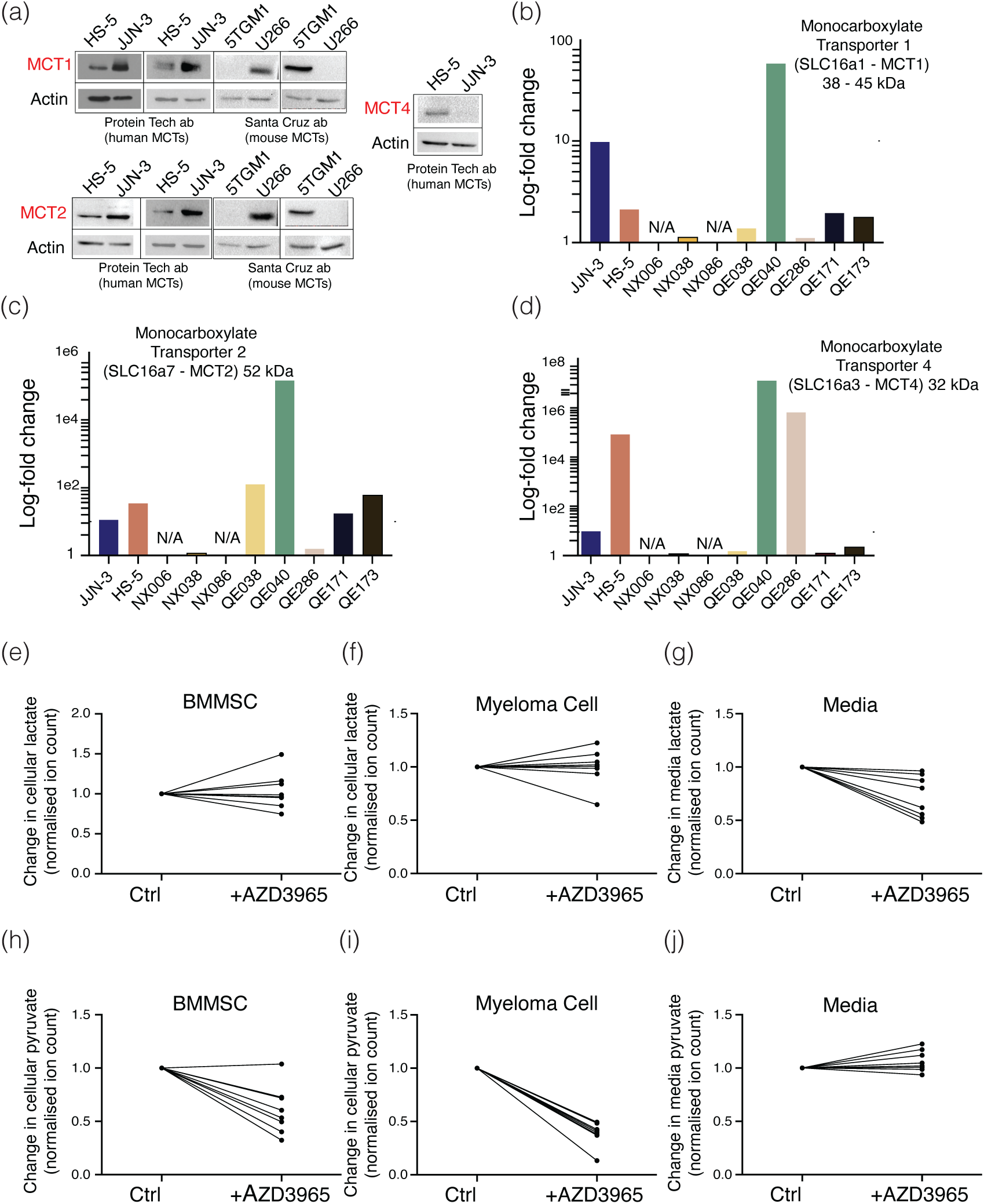
Expression and inhibition analysis of monocarboxylate transporters in HS-5 and JJN-3 cells. **(a).** Western blot analysis showing MCT-1 presence in both HS-5 and JJN-3 cell lines, as well as in control cell lines 5TGM-1 and U-266. **(b).** Western blot analysis showing MCT-2 expression across all tested cell lines. **(c).** Western blot analysis showing MCT-4 expression in HS-5 but not in JJN-3 cells. Actin is used as a loading control for all samples. **(d).** Quantitative PCR (qPCR) analysis of MCT-1 expression shows significant variability in log-fold changes across different patient samples, HS-5, and JJN-3 cell lines (see appendix). **(e).** qPCR analysis of MCT-2 expression levels across various cell lines. **(f).** qPCR analysis of MCT-4 expression, showing a notable peak in one patient sample and non-detectable levels in others. **(g-i).** Response of BMMSC and myeloma cells to MCT-1 inhibition by AZD3965. Lactate concentrations in BMMSCs and myeloma cells did not significantly change after AZD3965 treatment, with only a slight reduction in the media. **(j-l).** Experiments show a consistent decrease in pyruvate levels in both cell types and media after AZD3965 treatment.

We therefore expanded our analysis to primary myeloma and MGUS samples, and observed that transcripts for all three were expressed in both cell types (Figs. 2b to 2d), regardless of disease stage or other clinical readout (Appendix A). Of the three MCT transporters investigated, all have relatively high affinity for lactate, but of interest, MCT-1 and 2 have significantly higher affinity for pyruvate than lactate, contrasting with MCT-4 [60]. As pyruvate was predicted by the CBM to have net flux between the two cell types (see Vera-Siguenza et al. [24]), we therefore examined this interaction using a pharmacological inhibitor reported to inhibit MCT-1 with a Ki of 1.6 nM, and MCT-2 with a six-fold higher Ki [61]. To test the hypothesis that pyruvate is trafficked between BMMSCs and myeloma cells, we incubated primary autologous co-cultures from patients (Appendix A) with 10 µM AZD3965, a dose significantly in excess of the Ki for both MCT-1 and 2, and reported not to inhibit MCT-4 [61].

Our experiments revealed that across all donors, cellular lactate remained relatively unchanged in both BMMSCs and myeloma cells (Fig. 2e and 2f), with a small decrease observed in media lactate (Fig. 2g). Conversely, we observed a consistent decrease in pyruvate levels (Figs. 2h and 2i) across both cell types, which was particularly striking in the myeloma cells (Fig. 2j). Media levels were relatively unaltered, supporting the fact that MCT-1/2 facilitate bidirectional flux of their substrates. When the resulting change in the ratio of pyruvate-lactate was calculated from these studies, it clearly decreased in all three compartments, with the largest decrease apparent in the myeloma cells. These results support the hypothesis that treatment with AZD3965 has a more significant effect on intracellular pyruvate flux than lactate, which in turn alters intracellular pyruvate levels. The same was observed in JJN-3 cells when in co-culture with BMMSCs following treatment with AZD3965 (Fig. 3a).

**Figure 3:**
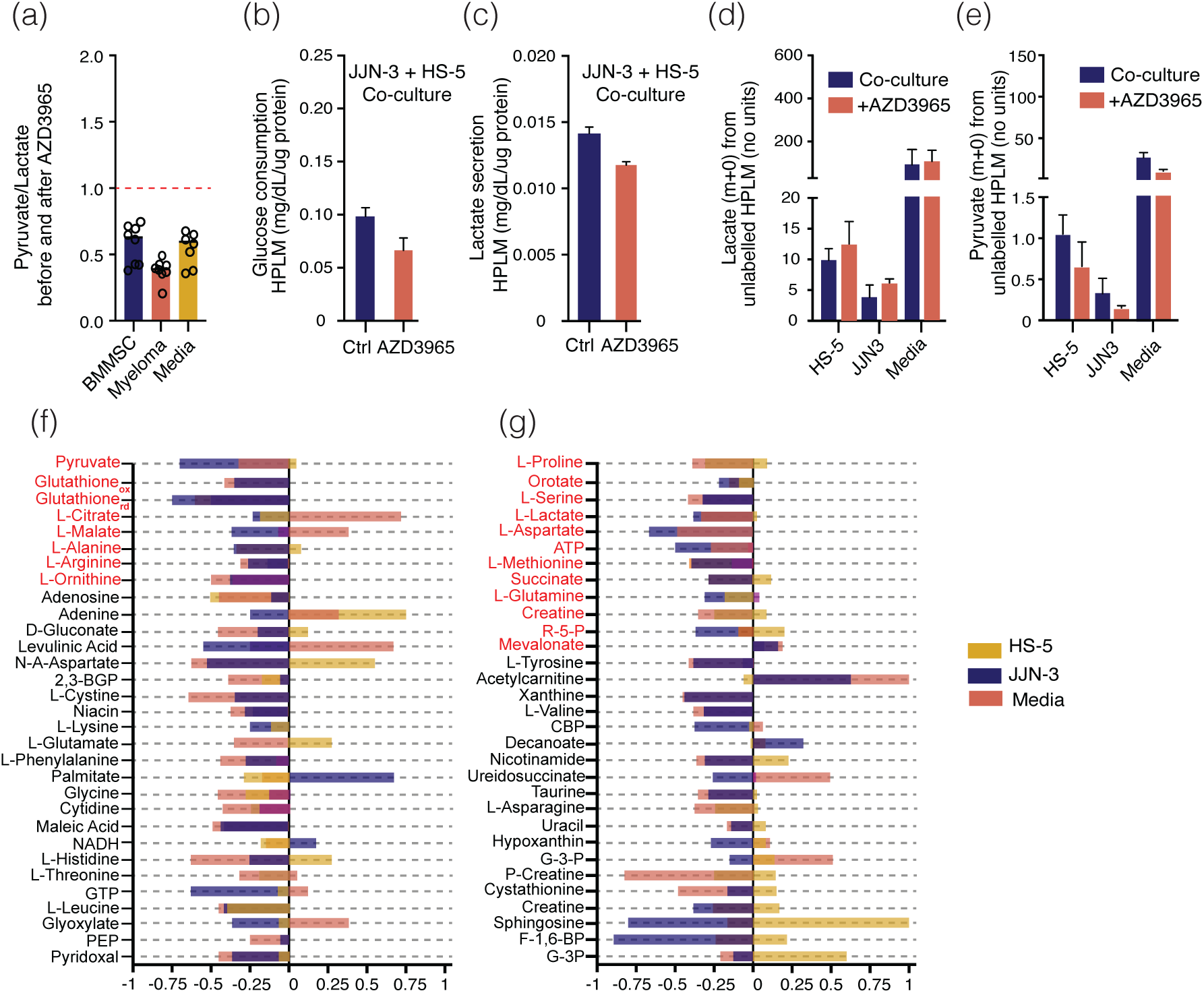
Comparative analysis of metabolic responses to AZD3965 treatment in BMMSC, myeloma cells, and media. **(a).** Comparative analysis of the pyruvate ratio before and after treatment with AZD3965 in BMMSC, myeloma cells, and media. **(b - e).** Impact of MCT-1 inhibition with AZD3965 on metabolic profiles using human plasma-like medium (HPLM) in co-culture. **(f** & **g).** Computational simulation results on MCT-1 deletion impact. Results show a reduced pyruvate flux in co-culture medium and myeloma cells. See Materials and Methods.

The directionality of metabolite flux facilitated by MCT transporters is a direct function of the relative concentrations of the metabolites on each side of the membrane. We therefore sought to validate our results in a medium that contained physiologically relevant lactate and pyruvate concentrations, and examined the effect of AZD3965-mediated inhibition of MCT-1/2 on pyruvate and lactate when used in Human Plasma Like Medium (HPLM). We observed that under these conditions, AZD3965 elicited a significant decrease in glucose consumption from the media, as well as a small decrease in lactate excretion (Fig. 3b and 3c) [62]. Intracellular lactate was observed to increase in both cell types, which alongside the decreased media lactate suggested that AZD3965 was inhibiting lactate efflux, albeit only by a small amount (Figs. 3c and 3d). However, a significant decrease in intracellular pyruvate was observed after treatment with AZD3965 (Fig. 3e), suggesting that pyruvate flux might be decreased, or alternatively increased reduction of pyruvate to lactate may also be occurring. These results suggest that inhibition of MCT-1 alters the intracellular redox ratio as previously reported, indicating a role for this transporter in maintaining redox homeostasis in myeloma cells.

We leveraged our in-silico CBM to simulate and screen a wider range of metabolic consequences upon inhibition of pyruvate influx/efflux through MCT-1/2 inhibition. These simulations were achieved by using the gene-rule deletion routine - delete-ModelGenes() - part of the MATLAB COBRA Toolbox, as detailed in our previous work and toolbox documentation [24, 40]. Our simulation results (Figs. 3f and 3g) indicated that the deletion of MCT-1/2-facilitated pyruvate transport led to a reduction in pyruvate flux, both in the culture medium and the myeloma cells. Note that this result is based on the extrapolated flux lower and upper bounds from our ^13^C-MFA experiments. These simulations also predicted an increase in the metabolism of intracellular pyruvate in the BMMSCs after inhibition of MCT-1/2, as the model predicted directionality of pyruvate flux from BMMSC to myeloma.

The model’s results are therefore supported by our experimental observations of pyruvate trafficking from BMMSCs to myeloma cells [24]. Beyond pyruvate, the CBM also indicates that the loss of MCT-1/2 activity has significant downstream effects on cellular carbon metabolism, predicting that intracellular pyruvate in myeloma is metabolised through the TCA cycle (Fig. 3g). Furthermore, predictions from the model – such as effects on nucleotide and lipid synthesis – are in line with our previously reported effect of reduced pyruvate uptake on myeloma cell biomass flux [24].

### Inhibition of MCT-1/2 with AZD3965 results in network-wide changes in myeloma cell metabolism

To biologically evaluate the modelling predictions, we conducted an untargeted metabolomic analysis on the cells and media from co-cultured JJN-3 and HS-5s in the presence and absence of AZD3965 (Supplementary Material Fig. 4). Through a differential analysis, we were able to observe that the vast majority of metabolic changes, upon MCT-1 inhibition, were decreases in abundance, suggesting that the transporter represents a central node in the regulation of the metabolic community. In agreement with our experimental observations, pyruvate, alanine, and lactate were decreased upon AZD3965 treatment (Supplementary Material Fig. 4). Our model predicted that loss of pyruvate carbons led to significant effects on a number of pathways that were required for anabolism, including oxidative TCA cycle activity. We therefore directly assessed this predicted change in metabolism by tracing the fate of ^13^C_6_-glucose after treatment with AZD3965. We observed that while relatively few significant changes in glucose metabolism were observed in HS-5 cells after AZD3965 (Fig. 4a), JJN-3 cells had reduced pyruvate levels, alongside aspartate and the related amino acid, asparagine. Increases in some TCA cycle metabolites were also observed (Fig. 4b and 4c).

**Figure 4:**
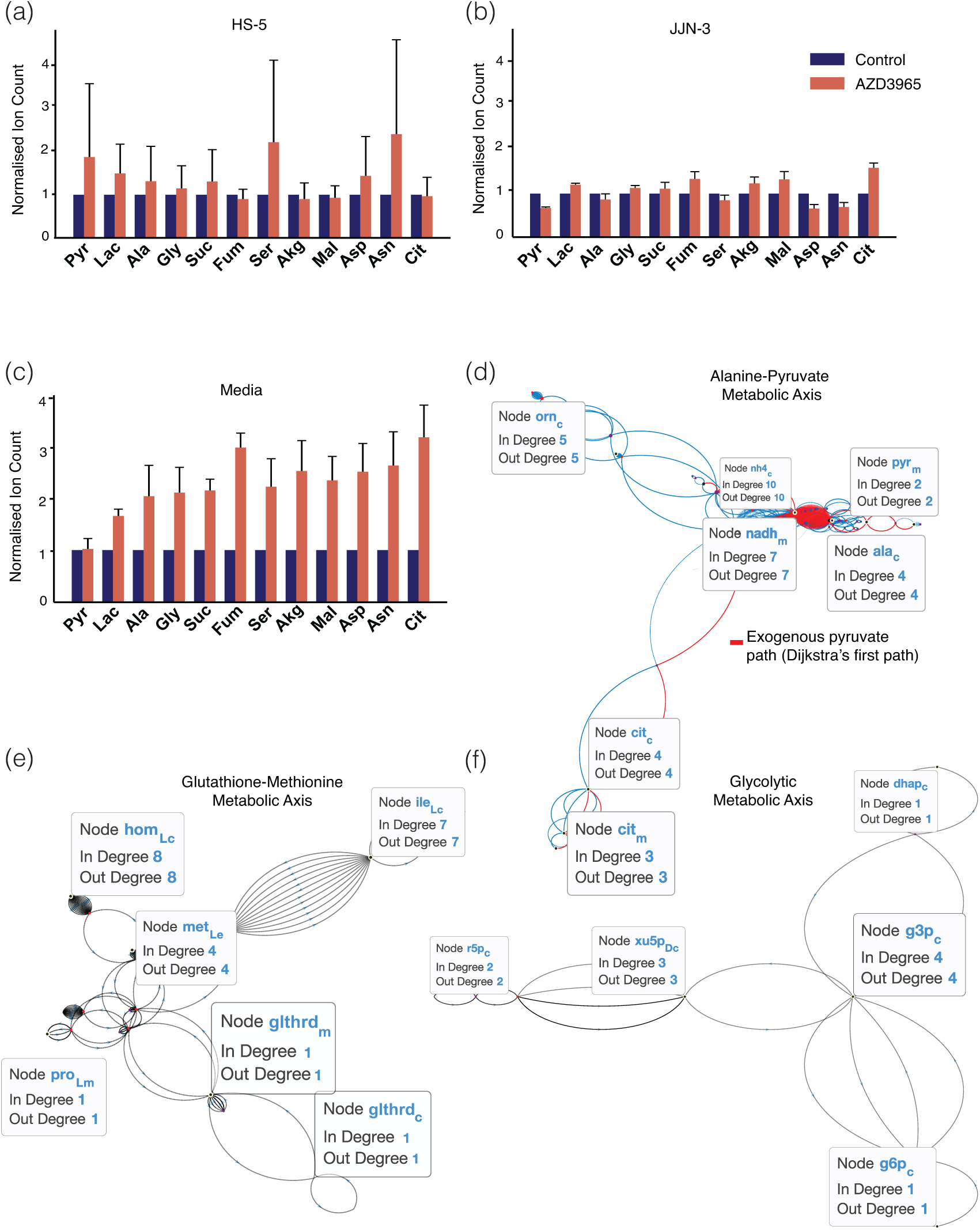
Metabolic impact of MCT-1 inhibition with AZD3965 on ^13^C_6_-glucose metabolism in co-culture, comparing HS-5 cells and JJN-3 cells to media control. **(a).** Displays minimal metabolic changes in HS-5 cells following AZD3965 treatment. **(b).** Shows JJN-3 cells exhibiting metabolic alterations. **(c).** Indicates important metabolic changes in the media. **(d-f).** Visual representation that translates the biochemical interactions of our myeloma cell CBM model into an expansive flux map using a directed graph approach Vera-Siguenza et al. [24]. This visualisation marks metabolites as nodes, and the transformations induced by reactions are represented as directed edges that represent the flow of metabolites along the distinct pathways. **(d).** Depicts the Alanine-Pyruvate metabolic axis, illustrating a significant pathway where pyruvate is a key metabolite, indicating an efficient pathway possibly due to alanine cycling. **(e).** Shows the Glutathione-Methionine metabolic axis, highlighting the connection between glutathione and methionine, which the model predicts influences redox regulation and antioxidant response. **(f).**Highlights the glycolytic metabolic axis, mapping the relationship between glycolysis and the pentose phosphate pathway, implicating a role in NADPH production and redox homeostasis. The size of each node (circle) represents the number of connections (degree), indicating the metabolite’s prominence in the cell’s metabolism. These metabolic axes underscore the interconnected nature of metabolic pathways and their potential alteration upon inhibition of pyruvate transport by MCT-1.

The ability to generate sufficient aspartate has previously been shown to be a central component of a cancer cell’s ability to support its proliferative drive, as the TCA cycle is bith driven and drives cellular NAD(P)(H) homeostasis. These data therefore corroborate our previously published model, which suggests that loss of pyruvate influx/efflux between the BMMSC and myeloma cell could reduce myeloma cell proliferation [13, 17, 24, 50, 63, 64].

### Shuttling of metabolites through MCT-1/2 supports malignant plasma cell redox balance

To further highlight other relationships between metabolite changes, we took our previously published model, and applied Dijkstra’s algorithm, to systematically examine the model’s metabolic network, starting from a specific metabolite – in this case pyruvate – and selected the nearest node, operating under the assumption of a natural tendency towards ‘minimal resistance’ [24, 65, 66, 67]. This analysis provided a visual dissection of the JJN3’s metabolome in co-culture with HS-5s (Fig. 4d-f). The results suggest the fate of pyruvate after import in JJN3 cells, as well as the effect of this on the wider metabolic network, can be collapsed into three critical ‘metabolic axes’. Each axis represents a key set of reactions, with its relative importance in the cell’s broader metabolic activity indicated by the number of encompassed metabolites (or nodes), represented by size. The primary axis (Fig. 4d) highlighted a clear link between alanine, pyruvate, and citrate, consistent with the alanine cycling hypothesis suggested in our previous work and experimentally described in other systems [24, 67, 68, 69, 70].

The secondary axis found in our model predicts (Fig. 4e) a linkage between glutathione, proline, and methionine, although this requires further experimental validation. Glutathione is central to the cellular antioxidant response, a metabolic network of which proline is also a significant member, while the amino acid methionine is involved in multiple pathways including 1-carbon metabolism and lipid headgroup synthesis. This node aligns with the metabolomic data suggesting increased uptake of glutathione into one or both cells in the co-culture system after MCT-1 inhibition, suggesting that pyruvate uptake into myeloma cells may impact wider redox regulation in the cells (Supplementary Material Fig. 4).

Finally, the third and smallest axis (Fig. 4f) suggests that inhibition of MCT-1/2 may alter the relationship between glycolysis and the pentose phosphate pathway – a major source of reducing potential (NADPH) required to respond to oxidative stress. Our model therefore predicts a significant shift in the control of redox homeostasis within the plasma cell through both axes 2 and 3.

### Exogenous glutathione supplementation partially reverses proliferative defect after AZD3965 treatment

Our investigations indicated that the loss of MCT-1/2 activity resulted in altered redox homeostasis and a metabolic profile that suggested decreased support for cellular anabolism. Interestingly, the most significant change observed in the media after inhibition of MCT-1/2 was in the antioxidant glutathione (GSH), suggesting that AZD3965 treatment incites a wider perturbation in cellular redox homeostasis (Figs. 5a). To investigate this experimentally, we treated co-cultured cells with AZD3965 and found around a 40% reduction in proliferation over the observed period (Fig. 5a). Given that our previous results suggested that myeloma cells may uptake glutathione under these conditions, we supplemented them with additional exogenous glutathione (GSH). We observed less label dilution in the media, indicating a reduced export into the media under AZD3965-treated conditions, and a partial recovery in proliferation after GSH addition, suggesting that indeed this could support continued proliferation of myeloma cells in the presence of AZD3965. Interestingly, a similar phenomenon has been observed in AML [12, 71]. Indeed, we observed increased uptake of GSH from the media, whether as a single moiety or after extracellular breakdown into its constituent (di)amino acids, in the presence of AZD3965 (Fig. 5b).

**Figure 5:**
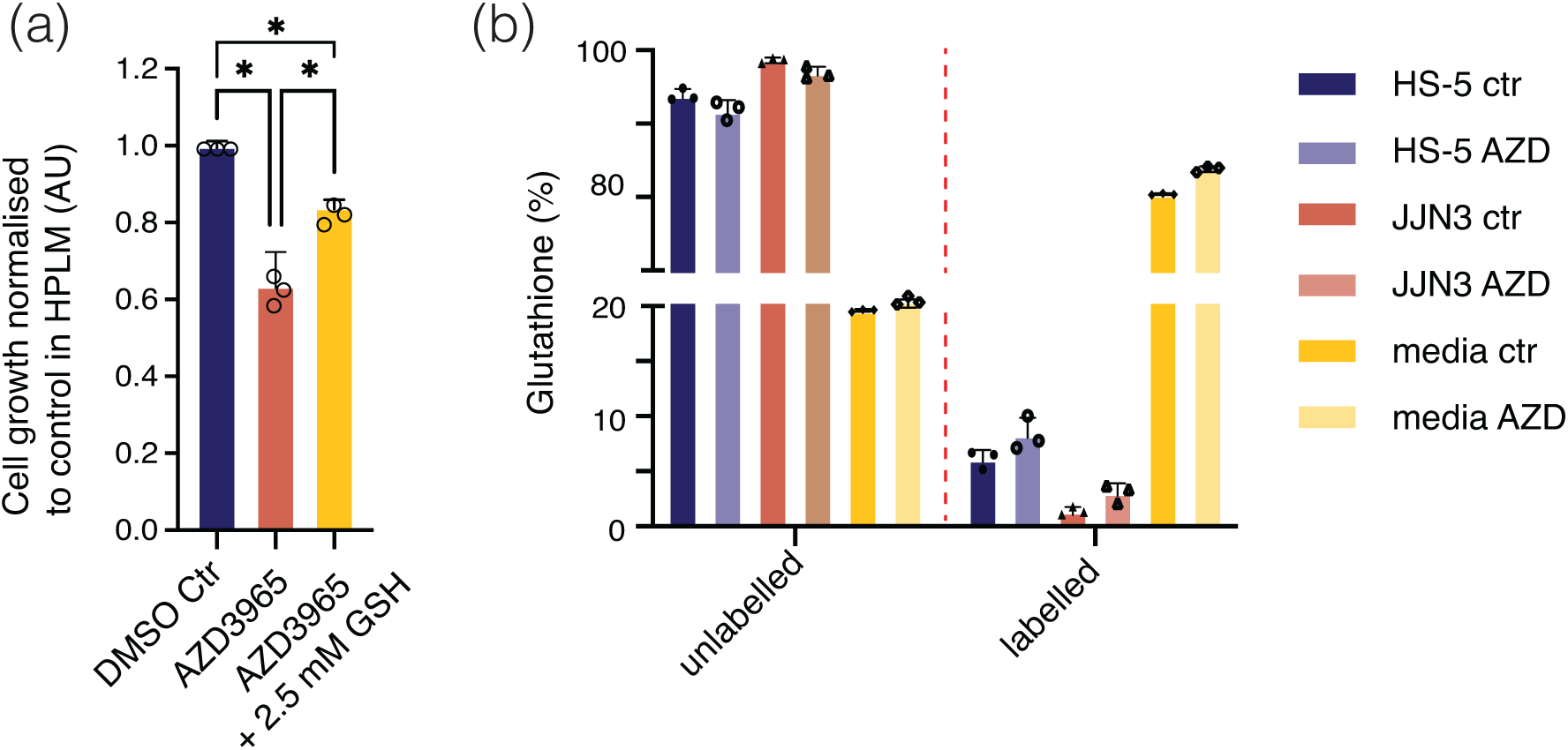
Metabolic responses to MCT-1/2 inhibition and glutathione supplementation in co-culture. **(a).** Effect of AZD3965 and GSH supplementation on JJN-3 cell proliferation - over 48 hours. The DMSO control is normalised to 1 on the y-axis. Treatment with AZD3965 leads to a decline in proliferation (approximately a 40% decline). Subsequent supplementation with 2.5 mM GSH partially restores proliferation. These results highlight a compensatory role by glutathione in response to MCT-1/2 inhibition **(b).** Changes in extracellular glutathione (GSH) enrichmment before and after MCT-1/2 inhibition with AZD3965 (right of dashed line). A reduction in media levels when treated with AZD3965 is indicative of an active GSH uptake by cells in response to reduced pyruvate import via MCT-1 inhibition with AZD3965. (*p*<*0.001 from ANOVA and Dunn’s multiple comparison).

## 4. Discussion

In this study, we investigated the metabolic interactions between myeloma cells and bone marrow mesenchymal stem cells (BMMSCs). Our study supports previous observations that suggest cancer-stromal metabolic interactions are not merely incidental but ‘cooperative’ [9, 10, 12, 17, 22, 24, 72]. To date, a number of studies have reported metabolic features of myeloma cells cultured in isolation [73, 74, 75, 76, 77]. In this investigation, we sought to expand on previous findings through an approach that utilised a co-culture system of BMMSC and myeloma cell integrated with mathematical modelling techniques to move closer to a model that better represents the bone marrow tumour microenvironment.

We found that, consistent with our earlier model predictions (Vera-Siguenza et al. [24]), co-culture of BMMSCs and myeloma cells permits metabolite sharing between the two. This setup supports the cancer cell metabolic network [24, 72, 78, 79, 80]. Our observations support a wider multicellular metabolic network in the bone marrow in which myeloma cells respond to their own metabolic tumour microenvironment, and also confirms the appropriateness of using constraint-based model to approach this computationally [24].

Our data suggest that the transport of monocarboxylates, including pyruvate and lactate, through MCT-1/2 is important for supporting myeloma cell proliferation. While MCT-4 has a higher affinity for lactate than for pyruvate, the reverse is true for MCT-1 and MCT-2, with pyruvate having a sub-millimolar affinity for MCT-1. Our model and experimental data, as illustrated in Figs. 3f and g, are consistent with myeloma cells taking up pyruvate from the media, using it to support multiple metabolic pathways both directly and indirectly. Although we have not formally excluded lactate from playing a role in this metabolic support, the presence of phosphorylated metabolites in the media, as predicted by the model, has not been reported for some of these metabolites and would need to be confirmed through experimental data. Despite this, our findings are in agreement with previous studies on metabolic interactions with cancer-associated fibroblasts (CAFs) [7, 9, 10, 11, 12, 22, 31].

Our findings do, however, contrast with previous findings in myeloma by Fujiwara et al. [9], Barbato et al. [10], Fujiwara et al. [22] and Walters et al. [81]; namely, MCT-1/2 being a primarlily lactate transporter in myeloma. However, we do agree in the clear role of MCT-1/2 in maintenance of a wider metabolic network. Through using Dijkstra’s algorithm, we created a detailed visualisation of the myeloma cell metabolome, which suggested three ‘metabolic axes’ predicted to have significant relevance in the metabolic network (Fig. 4) [65, 66, 67]. It highlighted a primary axis that included alanine, pyruvate, and citrate (Fig. 4d), alongside [11, 24, 82, 68, 83]. However, it also highlighted a second axis connecting glutathione (GSH), proline, and methionine. This latter axis was interesting given the role of GSH as a cellular antioxidant [12]. A role for glutathione was supported by metabolomic data (Supplementary Figure 4), as well as the partial result of the proliferative defect elicited by AZD3965 inhibitor treatment (Fig. 5a). Interestingly, our results agree with those from studies of AML, in which increased GSH was observed in AML cells co-cultured with BMMSCs [12, 84, 76, 85].

In summary, this study presents compelling preliminary data suggesting that AZD3965 treatment in patients with multiple myeloma may synergise with therapies targeting the cellular antioxidant response. However, further research is needed to determine the most effective combination, as it would likely need to be found outside of existing routine regimens [86, 87].

## Acknowledgements

Funding from Cancer Research UK to (DAT, EVS, and CEG) (C42109/A26982 and C42109/A24747). (AB) is supported by Cancer Research UK [SEBSTF-2021/100002]. (IC) is supported by a María de Maeztu Post-doctoral Fellowship. (FS) would like to acknowledge the funding for a UKRI Future Leaders Fellowship (MR/T043571/1). (CS) is supported by a Myeloma UK project award (MUK2021.ED05) to (CB, MD). The funders had no role in study design, data collection and analysis, decision to publish, or preparation of the manuscript.

## 5. Data Availability

The simulations carried out in this study were performed using the MATLAB 2021a software. Flux balance analysis, thermodynamic flux balance analysis, and flux variability analysis simulations were performed using MATLAB’s constraint-based reconstruction analysis (COBRA) toolbox in conjunction with the IBM optimisation CPLEX 12.7.1.package and Gurobi. ^13^C-MFA simulations and Monte Carlo parameter estimation were performed using the MATLAB-based graphical user interface isotopomer network compartmental analysis (INCA) suite software under an academic licence issued by Vanderbilt University. The supplementary material accompanying this article includes several generated scripts that are needed to run legacy algorithms, such as mCADRE with CPLEX and MATLAB 2021a. However, we have also generated a GitHub repository under the APACHE 2.0 license, where our model files can be found at: https://github.com/esig526/MetabolicNetworks

## Appendix A. Patient Data

| No. Sample | Date | Sex | Age | Disease Status | B2M (mg/L) |
| --- | --- | --- | --- | --- | --- |
| QE288 | 23/02/2023 | Male | 75 | Myeloma On First Line | 3.2 |
| QE293 | 03/04/2023 | Male | 61 | Multiple Myeloma Diagnosis | 14 |
| QE309 | 07/06/2023 | Female | 54 | Multiple Myeloma Diagnosis | 6.4 |
| QE315 | 27/06/2023 | Female | 72 | Multiple Myeloma Diagnosis | 2.1 |
| QE316 | 30/06/2023 | Male | 71 | Myeloma On First Line | 3.8 |

| No. Sample | Albumin (g/L) | ISS | PP Type | PP (grams/Litre) | % PC (Histology) |
| --- | --- | --- | --- | --- | --- |
| QE288 | 2 | 33 | IgG Lambda | <7.51 | $\leq 1$ |
| QE293 | 3 | 22 | IgG Kappa | 91.5 | 75 |
| QE309 | 3 | 41 | Kappa Light-Chain | < 0.01 | 30 |
| QE315 | 1 | 43 | IgG Lambda | 10.9 | 60 |
| QE316 | 2 | 40 | IgG Kappa | 3.8 | $\leq 5$ |

| No. Sample | FLC Ratio | Creatinine ($\mu\text{mol/L}$) | % PC (Flow) | % PC (Morphology) |
| --- | --- | --- | --- | --- |
| QE288 | 1.3 | 94 | $\leq 0.5$ | 3 |
| QE293 | 165.902 | 86 | 2 | 4 |
| QE309 | 391.51 | 143 | 7.2 | 10 |
| QE315 | 0.01 | 62 | 5.8 | 12 |
| QE316 | 1.28 | 93 | $\leq 0.2$ | 6 |

## Appendix B. Dijkstra’s Algorithm

When modelling metabolic networks, represented by flux maps from optimising CBMs, Dijkstra’s algorithm can be adapted to find the shortest paths in terms of metabolic fluxes between metabolites [65, 66]. Constraint-based models contain information regarding the stoichiometry of the reactions ocurring throughout the entire metabolic network, reaction bounds, as well as an objective function to be optimised. Dijkstra’s algorithm can be applicable to metabolic CBMs as their solutions (that is, the optimised flux map) can be represented as directed graphs where nodes represent metabolites, and edges represent the fluxes or reactions connecting these metabolites [65]. Here, the ‘shortest path’ in this context would represent the sequence of reactions that results in the least amount of ‘resistance’, or the most efficient pathway, from a starting metabolite to all others in the network [88]. Briefly, Dijkstra’s algorithm, applied to a metabolic network (or CBM) follows the following principles:

1. The adjacency matrix, which indicates the connections (fluxes) between metabolites (nodes), is transformed into another matrix *M^′^*. In this matrix, each non-zero entry *m_ij_* from the original matrix *M* is replaced by 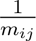 to indicate the ‘closeness’ or strength of the interaction between two metabolites. If *m_ij_* = 0, indicating no direct flux between metabolites *i* and *j*, then 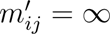 (is set to infinity), representing no direct pathway.
2. A starting metabolite is selected, analogous to selecting a start gene in genetic networks. This starting metabolite’s node is given a distance value of zero, while all other metabolites are assigned a temporary upper-bound value indicating they have not yet been reached by the algorithm.
3. The algorithm iterates through the network, selecting the metabolite (node) that can be reached with the minimal value from the start metabolite, updating the distances and creating a predecessor list, which tracks the pathway taken to reach each metabolite.
4. The process continues until all metabolites reachable from the start metabolite have been ‘visited’ and have a permanent distance value.
5. The output includes the shortest path distances to all reachable metabolites and the predecessor list that outlines the most efficient pathway taken.

This approach can reveal key metabolic pathways and identify ‘bottlenecks’ or critical reactions in the metabolism. Altering the starting metabolite or removing a metabolite from the network can significantly change the outcome of the analysis, highlighting the important role of certain metabolites in maintaining the metabolic network’s connectivity and efficiency. For implementation using a COBRA model we have uploaded, to our Github repository the script used to generate Figs. 4d to 4f.

