## Supplementary Material for "Pyruvate from bone marrow mesenchymal stem cells supports myeloma redox homeostasis and anabolism"

Elías Vera-Sigüenza†a,b, Cristina Escribano-Gonzalez†a, Irene Serrano-Gonzaloc, Kattri-Liis Esklad, Charlotte Speakmane, Alejandro Huerta-Uribe^h^, Lisa Vettorea, Himani Ranaa, Adam Boufersaouia, Hans Vellamad, Ramin Nashiebib, Ielyaas Cloete^i^, Jennie Robertsa, Supratik Basuf, Mark Draysone, Christopher Buncee, Guy Prattg, Fabian Spillb, Oliver D.K. Maddocks^h^ ,Daniel A. Tennanta

**Tennant (****).**

aInstitute of Metabolism and Systems Research, College of Medical and Dental Sciences, University of Birmingham, Birmingham, United Kingdom.

bSchool of Mathematics, University of Birmingham, Birmingham, United Kingdom.

cInstituto de Investigación Sanitaria Aragón, Fundación Española para el Estudio y Terapéutica de la Enfermedad de Gaucher y otras Lisosomales, Zaragoza, Spain.

dDepartment of Physiology, Institute of Biomedicine and Translational Medicine, University of Tartu, Tartu, Estonia.

eSchool of Biosciences, University of Birmingham, Birmingham, United Kingdom.

fRoyal Wolverhampton Hospitals, NHS Trust, Wolverhampton, United Kingdom.

gInstitute of Immunology and Immunotherapy, College of Medical and Dental Sciences, University of Birmingham, Birmingham, United Kingdom.

^h^School of Cancer Sciences, Wolfson Wohl Cancer Research Centre, University of Glasgow, Glasgow, United Kingdom.

^i^Centre de Recerca Matem\`atica. Edifici C. Campus de Bellaterra, 08193 Cerdanyola del Vall\`es, Barcelona, Spain

**Supplementary Material – Figures**

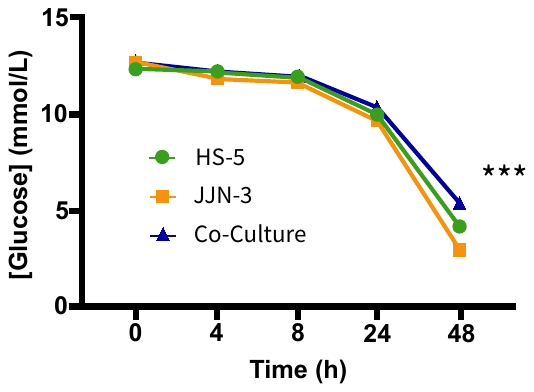

**Supplementary Fig.1:** Both, HS-5 and JJN-3 cells, bone marrow mesenchymal stem- and myeloma cells, respectively, consume less glucose in co-culture per cell than when cultured individually.

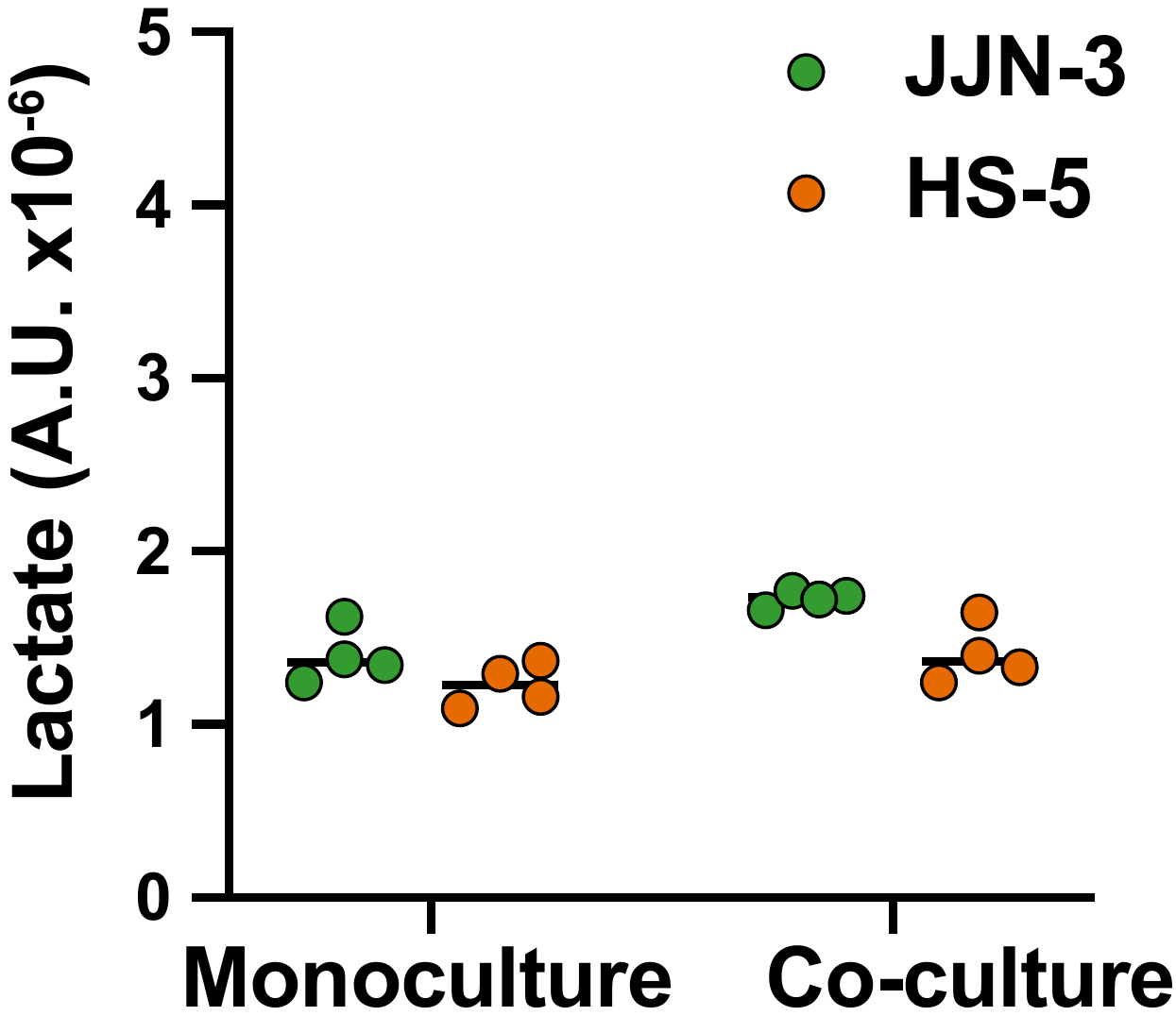

**Supplementary Fig.2:** The conversion of pyruvate to lactate in HS-5s showed no significant differences between monoculture and co-culture.

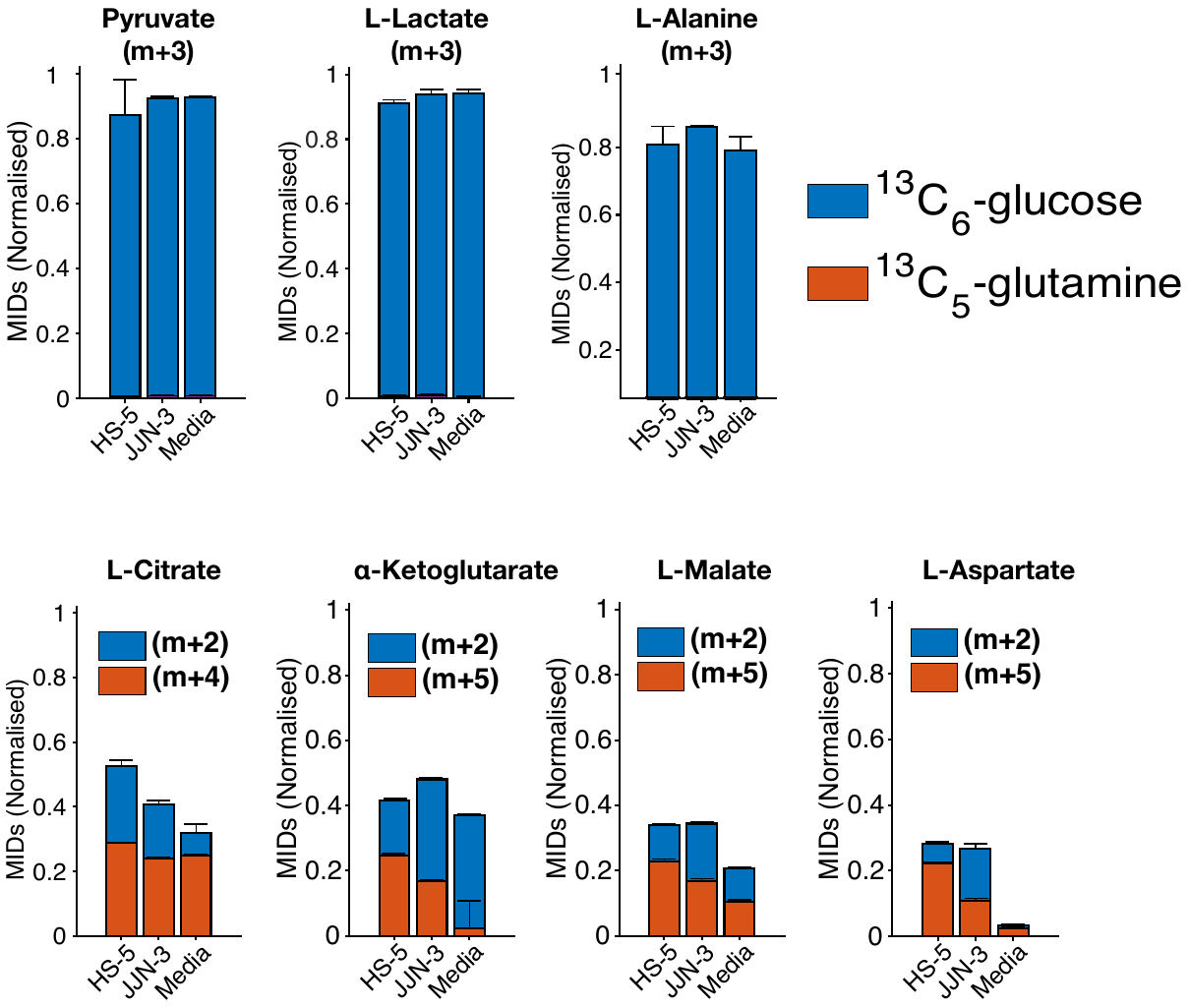

***Supplementary Fig.3:*** *Our experiment results imply the existence of an enhanced pyruvate oxidation process in both cell types when in co-culture.*

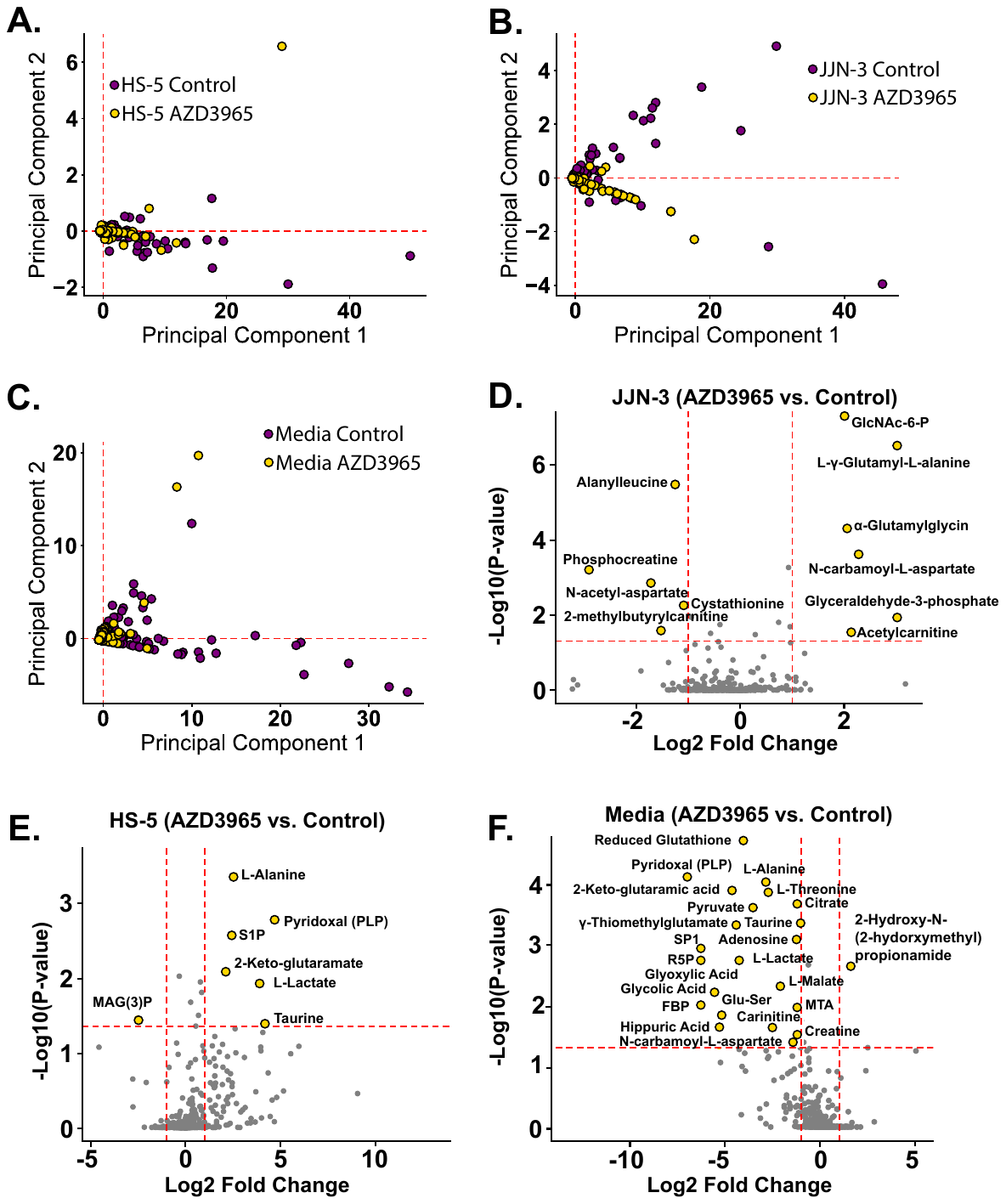

***Supplementary Fig.4:*** *In our study, we conducted an untargeted metabolomic analysis on co-cultured JJN-3 and HS-5 cells, both with and without the application of AZD3965, a potent inhibitor of the monocarboxylate transporter 1 (MCT-1). Our differential analysis revealed a predominant trend of decreased metabolite abundance upon MCT-1 inhibition, underscoring the critical role of this transporter in metabolic regulation within the cellular community. Consistent with our findings, key metabolites such as pyruvate, alanine, and lactate exhibited reduced levels following AZD3965 treatment, aligning perfectly with our experimental observations and highlighting the interconnectedness of metabolic pathways affected by MCT-1 activity.*
